# Efficient genetic editing of human intestinal organoids using ribonucleoprotein-based CRISPR

**DOI:** 10.1101/2023.03.31.535108

**Authors:** Nefeli Skoufou-Papoutsaki, Sam Adler, Paula D’Santos, Liz Mannion, Shenay Mehmed, Richard Kemp, Amy Smith, Francesca Perrone, Komal Nayak, Alasdair Russell, Matthias Zilbauer, Douglas J. Winton

## Abstract

Organoids are currently one of the most widely used *ex vivo* models in epithelial biology. Combined with genetic editing strategies, organoids offer a promise of rapid and efficient investigation of gene function in many models of human disease. However, to date, the editing efficiency of organoids with the use of non-viral electroporation methods has been only up to 30%, with implications for the subsequent need for selection including including turnaround time and exhaustion or adaptation of the organoid population. Here, we describe an efficient method of intestinal organoid editing using a Ribonucleoprotein CRISPR-based approach. Editing efficiencies of up to 98% in target genes were robustly achieved across different anatomical gut locations and developmental timepoints from multiple patient samples with no off-target editing. The method allowed us to study the effect of the loss of the tumour suppressor gene, *PTEN*, in normal human intestinal cells. Analysis of *PTEN* deficient organoids defined phenotypes that likely relate to its tumour suppressive function *in vivo*, such as a proliferative advantage and increased organoid budding. Transcriptional profiling revealed differential expression of genes in pathways commonly known to be associated with *PTEN* loss including mTORC1 activation.

## Introduction

Epithelial cells isolated from their resident tissue can be instructed to form three-dimensional self-organising and renewing organoid structures when placed in 3D matrices and provided with the appropriate niche signals (Sato *et al*., 2009). Organoids have bridged the gap between *in vivo* tissue based observations in human and mouse and those made in cell lines cultured in 2D. They have diverse applications from mechanistic studies of tissue development, to disease modelling and regenerative medicine (Rossi, Manfrin and Lutolf, 2018).

Organoid research has matured partly due to the concurrent development of technologies for genetic engineering, enabling studies of gene function (Fujii *et al*., 2015; Fessler *et al*., 2016; Kraiczy *et al*., 2017; van Rijn *et al*., 2018). Edited organoids have also been used for xenotransplantation approaches (Drost *et al*., 2015; Matano *et al*., 2015; Fumagalli *et al*., 2017). There are two main methods for introducing genome editing components into the cells; viral (retrovirus, lentivirus) and non-viral (electroporation and lipofection) (Teriyapirom, Batista-Rocha and Koo, 2021). Viral methods could traditionally reach efficiencies of 30-50% but more recent modifications of the transduction method have achieved efficiencies of 80-100% (Pirona *et al*., 2020; Gu *et al*., 2022). However, viral methods have associated biosafety and permanent integration issues while non-viral methods are simpler, minimising the time required for the production of the viral particles which can take up to 2-3 months (Lin *et al*., 2022). For the non-viral methods, electroporation has been found to result in higher organoid editing efficiencies (Fujii *et al*., 2015).

There are two approaches for genetic editing reagents based around delivery of either encoding vectors or ribonucleoprotein (RNP) (Figure 1a). For the RNP based approach a synthetic guide RNA is complexed with a purified recombinant Cas9 protein. This approach eliminates the need to translate the Cas9 protein, and has reduced off-target effects (Kim *et al*., 2014). More specifically, a 2.5-28-fold reduction in off-target effects has been reported with a RNP method compared to plasmid-based methods (Liang *et al*., 2015; Zhang *et al*., 2021). For these reasons, RNP based approaches are being rapidly adopted by the genome editing community (DeWitt, Corn and Carroll, 2017). However, so far, the vector-based approaches that have been used for editing of human organoids have resulted in efficiencies of only 10-30% (Fujii *et al*., 2015). This requires subsequent selection of the edited population, using either antibiotics or growth-factors and increases the time required for the generation of engineered populations and risks their exhaustion or adaptation during expansion.

**Figure 1:**
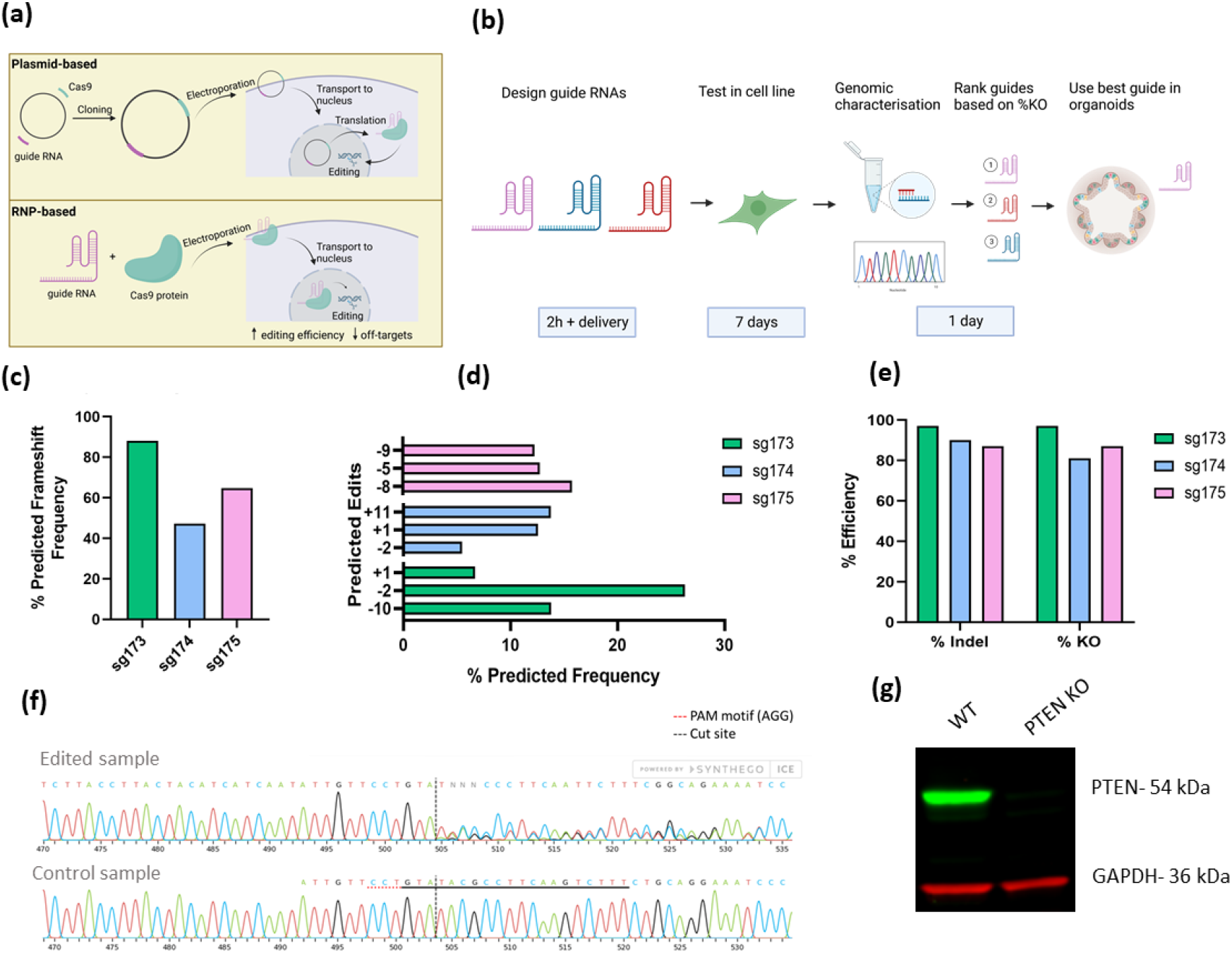
Experimental approach and optimisation of guides in cells. (a) Plasmid and ribonucleoprotein (RNP) approach for CRISPR-Cas9 genome editing. (b) Experimental approach for selection of best performing guide RNA in organoids and timeline. (c) Expected frameshift frequency of three PTEN guide RNAs. (d) Top three predicted edits for three PTEN guide RNAs. (e) Testing of three PTEN guide RNAs in MDA-MB-231 cells. % Indel includes all edits and % KO indicates edits that will lead to a frameshift. (f) Sanger trace for edited sample and control. Black dotted line indicates cut site and red dotted line indicates PAM motif. (g) Western blot for PTEN on PTEN WT and KO cells.

Here, we describe an efficient method for the generation of genetic knock-out human intestinal organoids using an RNP based CRISPR approach with the use of electroporation to deliver the editing components. Due to the high editing efficiencies, no subsequent clone selection is required. The method was applied to study the effect of the loss of the tumour suppressor gene, *PTEN*. Molecular and phenotypic characterisation of the edited organoids functionally validated the approach, and allowed behaviours likely relating to the tumour suppressive role of *PTEN* to be described.

## Results

### Generation and validation of *PTEN* KO organoids

For RNP mediated editing, a custom-made synthetic guide RNA is complexed with a recombinant Cas9 protein (Figure 1a). Guide RNAs were designed using online tools Benchling and Indelphi with a predicted On-target score >40 (Doench *et al*., 2016), Off-target score >80 (Hsu *et al*., 2013) and a frameshifting score of >80 (Shen *et al*., 2018). Three guides were designed per target gene to disrupt an early exon. Guides were first tested in a cell line to identify those with the best editing efficiency (% KO) as inferred following the deconvolution of Sanger sequencing using the <u>I</u>nference of <u>C</u>RISPR <u>E</u>dits software (Conant *et al*., 2022) (Figure 1b).

*PTEN* was selected as an initial target gene for disruption to generate a knock-out organoid model. It is an important tumour suppressor gene mutated in 6% of colorectal cancers (CRC) and germline mutation of which is causative for PTEN hamartoma tumour syndrome (Hobert and Eng, 2009; Cerami *et al*., 2012). The *PTEN* coding region is 8515 bp and the gene is located on chromosome 10q23.3 (Cunningham *et al*., 2022). The guide RNAs were designed to target exon 2 of *PTEN*. Of note, *PTEN* contains a highly conserved pseudogene, *PTENP1*, located at chromosome 9p21, with 98% sequence homology with the coding region of functional *PTEN* (Dahia *et al*., 1998). Based on the online design tool *PTENP1* is also predicted to be targeted with the designed guide RNAs due to this high sequence homology. When tested in cells, high editing efficiencies of up to 97% were seen with one of PTEN guides (Figure 1c-f). Loss of PTEN protein expression was confirmed in the edited cells (Figure 1g).

The best performing guide identified in cells was tested in human intestinal organoids. Different electroporation conditions, such as the commercially provided electroporation programs (see Methods), number of cells and varying Cas9 and guide RNA concentrations comprising the targeting complex, were tested to identify the combination leading to the highest editing efficiency (Figure 2a-b). The optimal condition was the use of 100,000 cells with the DS-138 program. Similar editing efficiencies of around 95% KO were detected with the complex concentrations for this program and the lower concentration, 5 μg:100 pmol, was used subsequently. Efficiencies of up to 98% were seen in three different patients and gut segments, duodenum, terminal ileum and sigmoid colon and remained stable over three weeks in culture (Figure 2c). High editing efficiencies were also observed in colonic organoids of a different developmental stage, that is human foetal organoids (Figure 2d).

**Figure 2:**
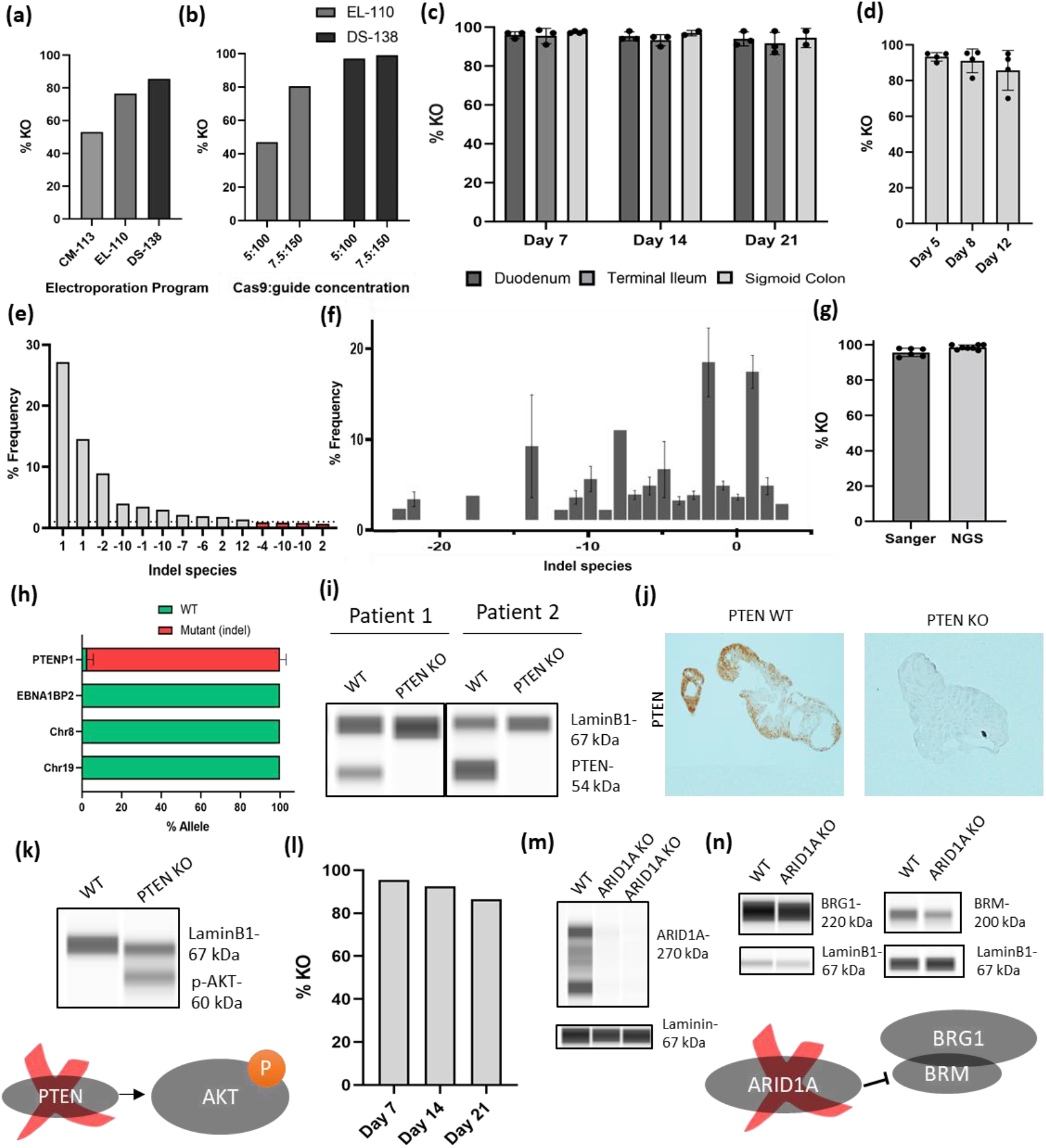
Efficient editing of human intestinal organoids. (a) Testing of different Lonza electroporation programs using 50,000 cells. (b) Two most efficient programs, EL-110 and DS-138 tested on 100,000 cells with 7.5 μg and 150 pmol or 5 μg and 100 pmol Cas9:guide RNA concentration. (c) PTEN electroporation using 100,000 cells, DS-138 program and lower Cas9 concentration on three different paediatric patients and three different gut segments. Percentage KO monitored over three passages. (d) Foetal organoid PTEN edited using same conditions as c for four patients. (e) Example of allelic species for one sample used for next generation sequencing (NGS). Dotted line indicates 1% allelic frequency, which was the set cut-off. Red bars indicate allelic species below the cutoff used. No WT reads were called. (f) Percentage frequency of different allelic species within all the PTEN KO samples submitted for NGS. Data from three patients and three timepoints. Includes species with >1% allele frequency. (g) Percentage KO assessed using NGS and sanger sequencing. (h) Off-target effects of PTEN guide assessed using NGS. Percentage WT (same size as predicted product) or mutant allele (contains insertions or deletions leading to a different sized product). Data from three patients. Off-target genomic coordinates: PTENP1: chr8:140155102, chr19:55504388. (i) Output of Wes (Biotechne) for PTEN protein on PTEN WT and KO organoids. (j) Immunohistochemistry for PTEN on PTEN WT and KO organoid FFPE sections. (k) Output of Wes (Biotechne) for p-AKT (ser473) on PTEN WT and KO organoids. (l) Organoid electroporation using ARID1A guide. Percentage KO monitored over three passages. Data from three technical replicates from one patient. (m) Output of Wes (Biotechne) for ARID1A on ARID1A WT and KO organoids. Data from two technical replicates. (n) Output of Wes (Biotechne) for BRG1 and BRM on ARID1A WT and KO organoids. Graphs show mean +STDV.

For validation of the genetic KO, samples were submitted for next-generation sequencing (NGS) covering the area targeted by the guide RNA. For the NGS analysis all the allelic species with a frequency higher than 1% were considered (Figure 2e). Species with indels leading to a different product size than the WT allele were considered mutant and were used to calculate the KO score. Even with this more sensitive sequencing method, the frequency of WT alleles across the edited samples was only up to 3% (Figure 2f). There was good correlation between the KO score estimated from the sanger sequencing and NGS (Figure 2g). In addition, primers were designed to cover the top regions predicted to have a potential off-target binding of the guide RNA based on the guide design tool (Benchling). These were composed of the top 3 off-target areas, including the *PTEN* pseudogene which has 100% sequence similarity with the *PTEN* guide RNA and two intergenic regions, in chromosomes 8 and 19. In addition, the highest off-target area within a gene, *EBNA1BP2*, was included. The anticipated editing of the PTEN pseudogene was observed, but none of the other areas were targeted by the guide RNA, with all the reads being of the WT sequence (Figure 2h). This shows that the method is specifically targeting the areas in the genome with the exact guide RNA sequence.

Next, the effect of the PTEN loss at the protein level was assessed. Loss of PTEN protein expression was observed using Wes™ (Biotechne) automated capillary based immunodetection and by immunohistochemistry (Figure 2i-j). A downstream target of PTEN, p-AKT, previously described as increased upon *PTEN* loss (Takao *et al*., 2018), was confirmed to be elevated in *PTEN* KO organoids (Figure 2k).

### Independent validation of organoid editing by targeting *ARID1A*

Electroporation conditions were applied with a different guide, targeting another tumour suppressor gene, *ARID1A*. High levels of KO efficiency were also observed (Figure 2l). Loss of ARID1A (SMARCA4) protein expression was confirmed (Figure 2m). SMARCA4 acts to regulate ATPases (BRM and BRG1) of the large ATP-dependent SWI/SNF chromatin-remodelling complexes involved in transcriptional regulation of gene expression and that are present in mutually exclusive complexes. To establish if there were downstream consequences to loss of ARID1A the expression of both ATPases was investigated. Of note, protein levels of BRM but not BRG1 were reduced demonstrating a functional consequence of *ARID1A* loss (Figure 2n).

These findings confirmed high KO efficiency with different gut segments, and organoids from different developmental timings and different guide RNAs. With the method described here, edited organoids can be generated within 26 days including the time required for adequate expansion of the culture and the design and testing of the synthetic guides (Figure 3).

**Figure 3:**
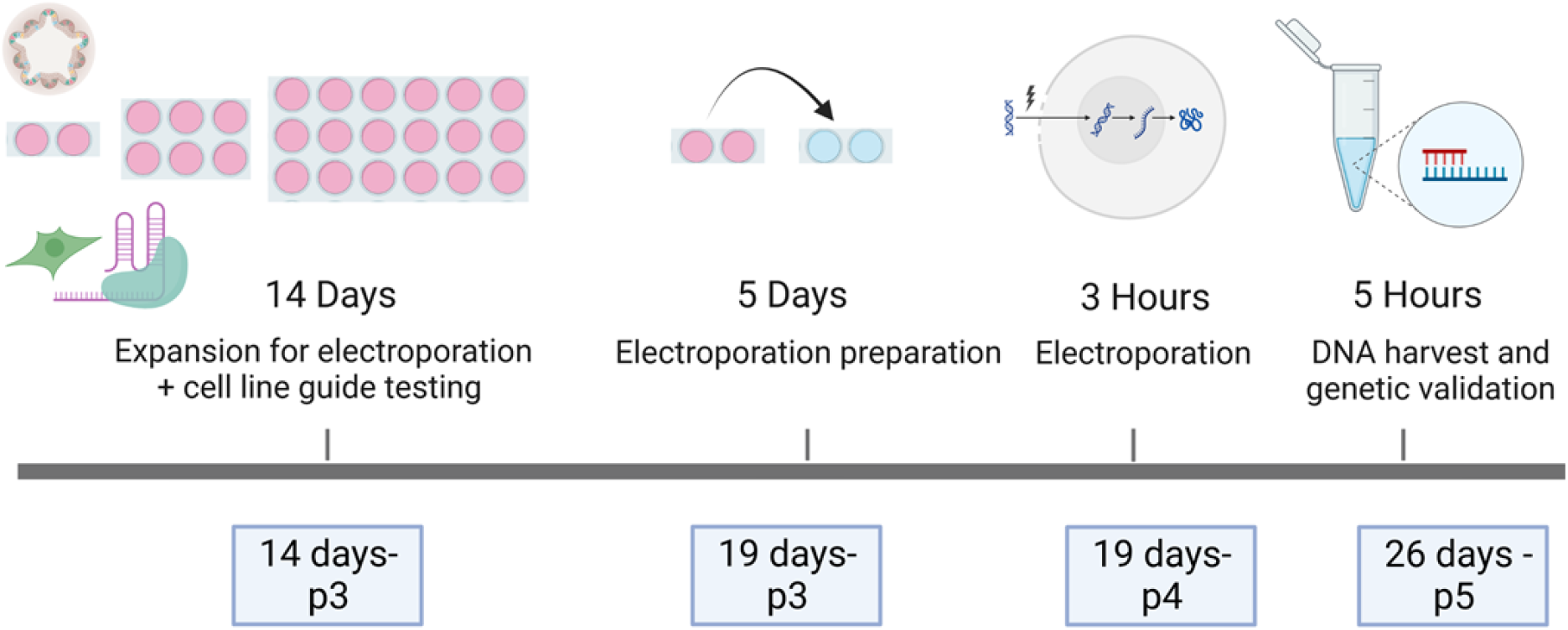
Outline of organoid electroporation process and timeline. Knock-out organoids can be generated within 26 days and 5 passages, starting from two wells of intestinal biopsies. This timing includes also the testing of target guide RNAs in cell lines.

### PTEN KO organoids exhibit increased budding and proliferative advantage

The phenotypic effect of *PTEN* loss in a human sigmoid colon organoid model was next assessed to explore how the normal intestinal cells might be affected in a way that could relate to cancer development, which can act as a functional validation of the model.

*PTEN* KO organoids appeared morphologically larger, with a higher cell number and exhibited an increased number of budding structures compared to WT organoids (Figure 4a-g). An increase in cell proliferation was also observed upon PTEN loss, as indicated by a higher proportion of cycling cells marked by S-phase marker MCM2 (Figure 4h-i).

**Figure 4:**
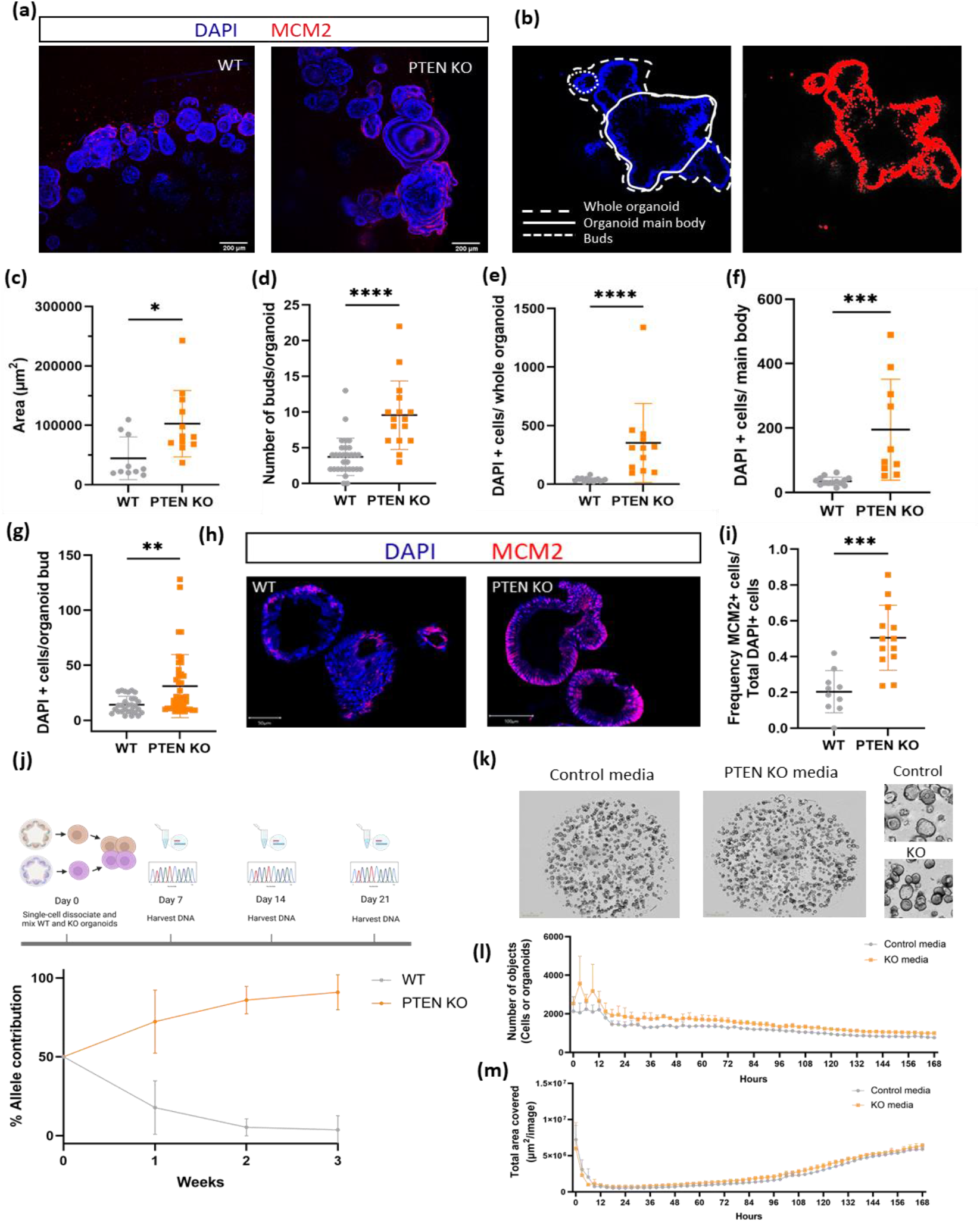
*PTEN* KO organoids show increased budding and exhibit proliferative advantage. (a) Wholemount staining of organoids. Shown as maximum projection. (b) Definition of organoid regions and counting of cells in Fiji. (c) Organoid area (μm^2^). Mann-Whitney test. p-value= 0.0112. (d) Number of buds per whole organoid. Mann-Whitney test. p-value < 0.0001. (e) Number of DAPI positive cells per whole organoid. Mann-Whitney test. p-value: < 0.0001. (f) Number of DAPI positive cells per main body. Unpaired t-test. p= 0.0009. (g) Number of DAPI positive cells per organoid bud. Mann-Whitney test. p-value= 0.0019. (h) Immunofluorescence MCM2 staining on FFPE organoid sections. c-g N= data from two patients and three technical replicates. (i) Frequency of MCM2 positive cells. N= data from three patients. Unpaired t-test. p= 0.0002. (j) Competition assay. Top schematic of experimental outline. Bottom WT and PTEN KO allele contribution over three weeks (21 days). N=data from three patients. (k) Images of WT organoids cultured with WT or PTEN KO conditioned media, 7 days post single-cell dissociation. Inlet showing top right corner of each well in higher magnification. (l) Quantification of organoid numbers in conditioned media experiment. (m) Quantification of organoid area in conditioned media experiment. N= data from three technical replicates. Graphs show mean +STDV.

A competition assay was set up to assess whether *PTEN* KO cells have a competitive advantage over WT cells in organoid cultures (Figure 4j). Indeed, after mixing *PTEN* KO cells with WT cells in a 50:50 ratio, the KO cells completely took over within three weeks in culture, suggesting either a direct proliferative advantage or possibly that they act to inhibit the growth of WT organoids. To investigate the latter WT cells were grown with *PTEN* KO conditioned media derived isogenically (Figure 4k-m). No effect on the WT organoid numbers or their size was observed, suggesting that PTEN KO cells outcompete WT cells due to a functional proliferative advantage.

### Transcriptomic characterisation of PTEN KO organoids

Bulk RNA-sequencing was performed to correlate the molecular changes with phenotypic alterations in *PTEN* KO sigmoid colon organoids. Three isogenic pairs of WT and PTEN KO patient derived organoids were profiled. Principal component analysis clustered organoids based on the originating patient rather than the genotype, suggesting a strong patient effect which was incorporated in the differential expression model (Figure 5a-b). There were in total 730 differentially expressed genes with 290 of them being upregulated and 440 being downregulated (False Discovery rate <0.05).

**Figure 5:**
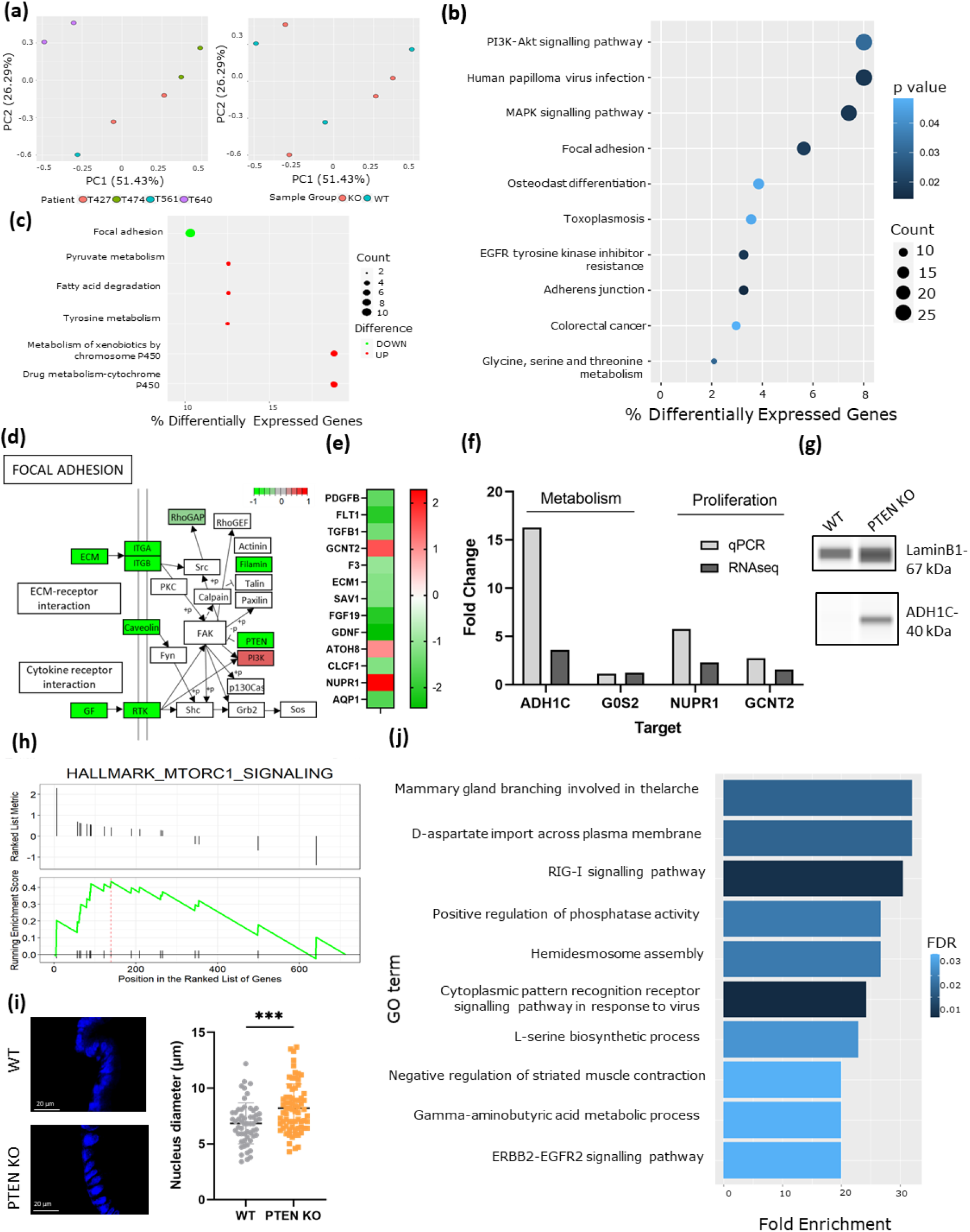
Transcriptomic profiling of *PTEN* KO organoids. (a) Principal component analysis based on patient ID. and genotype. (b) Annotated KEGG pathways for differentially expressed genes. (c) Annotated KEGG pathways for most highly differentially expressed genes (log Fold change > 0.7 and < −0.7). (d) Focal Adhesion pathway with highlighted genes being differentially expressed. Focus on the membrane side. (e) Regulation of cell proliferation identified as one of the gene ontology processes that is enriched in the dataset. Showing genes involved in regulation of cell proliferation with biggest fold change (>1 and <−1). (f) qPCR validation of fatty acid degradation targets ADH1C and G0S2 and proliferation regulators NUPR1 and GCNT2. Comparison with RNAseq fold change. Data from three patients. (g) Wes (Biotechne) protein validation of fatty acid target ADH1C. h) Gene Set Enrichment Analysis Hallmark Pathways. mTORC1 pathway (enrichment score: 0.44, pvalue 0.047). (i) Quantification of cell size. Cell diameter (μm). (j) Gene Set Enrichment Analysis of Gene Ontology terms based on top 500 differentially expressed genes ranked by ascending p-value. Showing top 10 Gene Ontology (GO) terms.

Pathway analysis of differentially expressed genes indicated enrichment for cancer-associated KEGG pathways such as the PI3K-AKT pathway, of which PTEN is a negative regulator, in addition to the MAPK signalling pathway and Colorectal cancer (Figure 5b). Changes in adhesion and adherens junctions were also highlighted. When focusing on the genes with the highest fold changes, focal adhesion was the top downregulated pathway in *PTEN* KO organoids (log FC< −0.7) while alternative metabolic pathways such as fatty acid degradation and tyrosine metabolism were included in the most highly upregulated pathways (log FC > 0.7) (Figure 5c). Several members of the focal adhesion pathway were downregulated, including cell receptor FLT1/VEGFR, or ligand PDGFB. Downregulation of collagens 9 and 6 (COL9A3 and COL6A1) was also observed. The majority of the downregulated genes encode proteins localised on the cell membrane (Figure 5d).

There was a general enrichment of the regulation of cell proliferation process, with some important genes such as *TGFB1* and *GCNT2* being differentially expressed (Figure 5e). One of the most highly upregulated genes in the PTEN KO dataset, *NUPR1*, is also involved in this process. Orthogonal validation of the most upregulated genes involved in the fatty acid degradation pathway, *ADH1C* and *G0S2* and proliferation regulatory genes, *NUPR1* and *GCNT2* was performed using qPCR (Figure 5f). The upregulation of fatty acid metabolism target ADH1C was also confirmed at the protein level (Figure 5g).

In addition, the mTORC1 pathway was found to be enriched in the *PTEN* KO organoids (Figure 5h). This pathway is an important regulator of cell proliferation and metabolism. mTORC1 is also known to be important for controlling the cell size, with activated mTORC1 leading to larger cells (Fingar *et al*., 2002). The size of a cell is known to be proportionate to the size of its nucleus (Edens *et al*., 2013) so the activation of mTORC1 in *PTEN* KO cells could result to a larger cell size and nucleus. The nucleus diameter of *PTEN* KO and WT organoids was measured and indeed, *PTEN* KO nuclei were found to be larger (Figure 5i), in line with the predicted effect of mTORC1 activation.

Finally, in terms of processes that might relate to the increased budding phenotype, enrichment of the mammary branching during breast development was implicated in the *PTEN* KO organoids with three of six genes (AREG, pHB2 and TGFB) known to be involved in the process being differentially expressed (Figure 5j).

## Discussion

Primary organoid cultures contain much of the diversity present in the donor and avoiding the need for selection following gene editing serves to minimise the time required for expansion (both before and after electroporation), and retains more of this biological diversity. The efficiency of the genome editing method described here was similar to viral delivery methods without the associated biohazard and mutagenesis issues. The use of the RNP method allowed an almost 3-fold increase the editing efficiency compared to previously published electroporation-based delivery methods using a plasmid-based approach (Fujii *et al*., 2015). The RNP-based CRISPR method was shown to be extremely robust, working with similar efficiencies for organoids taken from gut segments of different anatomical locations and developmental timepoints from multiple patients and for different target genes. Gene edits remain constant over time in culture and the editing was restricted to regions with 100% of sequence homology with the guide RNA. Although not tested, the method will likely translate to different epithelial organoid systems. A protocol for the RNP-based CRISPR approach, was recently published for the generation of FBXW7 KO intestinal organoids (Chan, Collins and Buczacki, 2023) and independently confirms the potential for improved editing efficiency in organoid models.

Functional and molecular analyses of edited organoids demonstrate the potential of the method to reveal biological insights. The generation of an *ARID1A* genetic KO demonstrated a reduction in the levels of one of the two downstream active mediators, BRM but not BRG1, which has not to our knowledge been previously identified.

The morphological and transcriptomic characterisation of the *PTEN* KO organoids, validated the genetic KO, with previously reported changes in signalling pathways and processes associated with loss of *PTEN* also seen in the organoid model. Loss of *PTEN* was associated with a proliferative advantage, metabolic rewiring and reduction in the process of focal adhesion. It was particularly interesting to note the two-fold downregulation of negative regulator of cell proliferation*, TGFB1*. Crosstalk of the TGFB and PTEN/PI3K/Akt pathway has been previously reported (Taniai *et al*., 2009). Moreover, it has been suggested that loss of PTEN may contribute to the differential role of TGFB signalling in the process of carcinogenesis, by promoting the tumour enhancer role of TGFB with specific effects on cellular motility and invasion (Hjelmeland *et al*., 2005). *TGFB1* has also been shown to influence the levels of PTEN protein expression in a prostate cancer model (Kimbrough-Allah, Millena and Khan, 2018).

The increase in cell proliferation in *PTEN* KO organoids could be fuelled by a metabolic switch regulated by the mTORC1 pathway. Activation of mTOR upon *PTEN* loss has been previously shown in different contexts including ovarian, head and neck squamous cell carcinoma and neurons (Matsumoto *et al*., 2016; Chui *et al*., 2019; Tariq *et al*., 2022).

*PTEN* loss also caused increased organoid budding. Components of the EGF and TGFB pathways are potentially implicated in this process by their involvement in branching morphogenesis. Activation of TGFB pathway has also been shown to inhibit ureteric duct growth while knockdown of TGFB was associated with an increase in duct budding (Clark, Young and Bertram, 2001; Nagalakshmi and Yu, 2015). Relatedly administration of EGF has been shown to reduce crypt fission in the colon (Bashir et al., 2003). Further work is required to determine if the increase in colonic organoid budding seen in *PTEN* KO organoids is mediated via altered TGFB or EGFR/AREG signalling.

To gain an insight into the process of cancer development, previous studies have attempted to replicate and test the order of gene mutation in colorectal cancer using organoids by sequential editing of the relevant events (Matano *et al*., 2015). For example, *SMAD4*, instead of *PTEN*, was modelled in that study as a late event. The method employed here to generate KO organoids without the need for selection changes that calculus. Any tumour suppressor gene can be mutated to determine its functional role at different stages of the neoplastic process. Illustratively, the involvement of PTEN in the focal adhesion pathway via direct regulation of focal adhesion kinase (FAK) has been reported (Tamura *et al*., 1998; M Tamura *et al*., 1999). These reports described a downregulation of focal adhesion formations in NIH 3T3 cells engineered to overexpress *PTEN* in apparent contradiction to the effect shown here where the same effect resulted from *PTEN* loss. This difference demonstrates the need to test gene, and cancer driver, functions in biologically relevant contexts such as those provided by primary organoids that can be efficiently gene edited.

## Methods

The in-detail description of the organoid electroporation using RNP CRISPR method protocol can be found here: DOI: dx.doi.org/10.17504/protocols.io.5qpvor74xv4o/v1 (Private link for reviewers: https://www.protocols.io/private/00104898A61E11EDA02B0A58A9FEAC02 to be removed before publication.)

### Human intestinal samples

Intestinal mucosal biopsies were collected from sigmoid colon, terminal ileum and duodenum from children under 16 years old undergoing diagnostic colonoscopy at Addenbrookes Cambridge Hospital. This study was conducted with informed patient and/or carer consent as appropriate, and with full ethical approval (REC-12/EE/0265). Foetal intestine was obtained with ethical approval (REC-96/085) and informed consent from elective terminations at 8–12-week gestational age.

### Human intestinal organoid culture and maintenance

Intestinal organoids were cultured in ADF+++ media referred here as WENRAFI (Suppl. Table 1). Media was replaced every 2-3 days. The standard organoid culture conditions were adapted from those previously described by replacing the p38 inhibitor with IGF-1 and FGF-2 (Fujii *et al*., 2018).

On passaging, organoids were disaggregated by intense pipetting. Following centrifugation (500g, 4minutes for all steps) the pellet was reseeded in fresh Matrigel (20 μl per well, BD356231) in 48 well plate with 250 μl of WENRAFI plus 10 μM Y-27632, ROCK inhibitor, for the first 24 hours.

### Design of synthetic guide RNAs and primers

Guide RNAs were ordered from Synthego (Suppl. Table 2). Three guide RNAs were designed per target using Benchling (Benchling [Biology Software]. (2023). Retrieved from https://benchling.com) and Indelphi (Shen *et al*., 2018) (https://indelphi.giffordlab.mit.edu/). Guides with a predicted high on target (>40), off-target (>80), and frameshift (>80%) score were selected preferentially targeting an early exon of the gene. Their knock-out efficiency was first assessed in cells. The guide yielding the highest knock-out per gene was taken forward for use in organoids.

Primers were designed on Primer 3 so that the target region is 150bp upstream and downstream from expected edit site to allow successful comparison of sanger sequencing traces, with products ranging from 500-800 bp (Suppl. Table 2).

### Electroporation related organoid culture

Organoids were passaged 5 days prior to the electroporation. Media was changed to reduced ENAFI medium (without Wnt and R-spondin) supplemented with 5 μM CHIR99021 and 10 μM Y-27632 (ENAFI-CY+) 48hours before electroporation and then changed to ENAFI-CY+ containing 1.25% (vol/vol) DMSO 24 hours before electroporation (Suppl. Table 3). For electroporation organoids were seeded in ENAFI-CY+ media with 1.25% (vol/vol) DMSO. The next day media was changed to WENRAFI + 10 μM Y-27632.

### Electroporation using Ribonucleoprotein Complex (RNP)

Parts of the protocol have been modified from Fujii *et al*., 2015 for RNP delivery in cells using electroporation.

On electroporation, the culture medium was removed and 500 μl of TrypLE Express supplemented with 10 μM Y-27632 were added to each well. Matrigel was scraped and the organoids were transferred into Falcon tubes and placed in a 37 °C water bath for 30 min with vigorous pipetting every 5 minutes achieve single cell dissociation. ADF basal medium was topped up to 10 ml and the tubes were centrifuged. True Cut Cas9 v2 protein (Thermofisher, A36499-500 ug at 5ug/ul) and full length, synthetic guide RNAs (previously reconstituted with water to 100pmol/ul) were thawed on ice. The supernatant was aspirated and the cells were re-suspended in 0.5 ml Opti-MEM medium. The number of cells was counted (unless otherwise stated 100,000 cells/condition were used). The RNP complex was made by mixing 1μl of 5 μg Cas9 and 1μl of 100pmol guide (1:3.3 ratio). The complex was incubated for 20 minutes at room temperature. A negative, no-Cas9, control and at least three technical replicates were always included. Cells were centrifuged, the supernatant aspirated and 300μl of PBS added. Cells were centrifuged and the supernatant was aspirated and pellet was resuspended in the appropriate volume of P3 supplement buffer (20μl per condition by mixing 16.4μl P3 buffer and 3.6μl of supplement 1, Lonza, V4XP-3032). The cells were transferred into 16-well nucleovette strips (Lonza). After 10 minutes at room temperature electroporation was performed on the Amaxa 4D Nucleofector (Lonza) with the DS-138 custom program unless otherwise stated. The cells were then incubated for 10 minutes at 37 °C and then 80μl of pre-warmed media was added to the electroporation chambers. Cells were centrifuged, supernatant discarded, and pellet was resuspended in 20μl of Matrigel per condition and seeded in 48-well plates (100,000 cells per well).

### Screening of gene edited organoids

One third of organoids in each well were harvested for screening at 7 days post-electroporation. DNA was extracted using the Arcturus PicoPure kit. Organoids were centrifuged and the pellet resuspended in 20μl of Proteinase K buffer from the Arcturus DNA PicoPure kit (ThermoFisher, KIT0103). They were incubated for 3 hours at 65°C followed by 10 minutes incubation at 95°C.

Genomic DNA was used for PCR amplification of the targeted exon in the gene of interest using the Q5 Polymerase enzyme according to manufacturer’s instructions (M0491, NEB), with 2 μl of organoid lysate in a 50 μl reaction. The product was cleaned using ZymoResearch PCR cleanup kit (D4013) and submitted for Sanger sequencing in both directions.

To quantify the a gene knock-out score, the Sanger sequencing traces were deconvoluted using the Inference of CRISPR Edits (ICE) tool (Conant *et al*., 2022).

### Targeted-amplicon next generation sequencing

DNA from three patients and two technical replicates, over three timepoints was subjected to targeted amplicon next generation sequencing. Regions of interest included the area targeted by the PTEN guide RNA, and the top off-target regions as predicted by Benchling. Primers were ordered with the addition of the CS1 and CS2 Illumina adapters. To amplify the regions of interest, PCR was performed using Phusion polymerase (NEB). Unique Fluidigm sample barcodes were added using Fast Start High Fidelity PCR System (Roche). Library was balanced based on the Bioanalyser molarity. Column-based Clean & Concentrator Kit (Zymoresearch) was used for PCR-cleanup. Primer dimers were eliminated by broad range (200-400 bp) size selection using the Blue Pippin (Sage Science) and samples were submitted for 150-bp paired end Miseq Nano Illumina sequencing.

For the analysis, paired end assembler for Illumina sequences, PANDAseq was used to merge corresponding forward and reverse reads into an *in silico* amplicon (Masella *et al*., 2012). The top 10 different species that started and ended with the correct primer sequence were identified, their frequency calculated and their size was noted. Based on the predicted amplicon size, they were then classified as mutant reads with indels or WT reads with no difference in the predicted product length.

### Organoid FFPE plug generation

Organoids were fixed in 4% PFA for 20 minutes at 4 °C. Giemsa dye was added (1:10 in 70% ethanol) for 1 hour. After washing with 70% ethanol, organoids were resuspended 2% pre-heated low melting point agarose and transferred on the cap of an Eppendorf, which was used as a mould. The plug was removed and added in 10% NBF overnight at 4 °C. Plugs were embedded in paraffin and cut into 5 um sections.

### Immunodetection

Antigen retrieval was performed using citrate buffer (10 mM sodium citrate, pH=6) with a laboratory pressure cooker (125 ° C for 2.5 minutes and then 91 ° C for 10 seconds). Blocking followed with 10% donkey or goat serum (Dako) or 3% Bovine Serum Albumin (BSA) incubated for 30 minutes. Primary antibody was incubated overnight at 4°C (Suppl. Table 4). The next day, secondary antibody (JacksonImmunoresearch biotin-SP-conjugated AffiniPure donkey anti-mouse or anti-rabbit, AB_2340785 and AB_2340593, 1:500 dilution for immunohistochemistry and Alexa fluorophore conjugated antibody, 1:200 for immunofluorescence) was added for 40 minutes (with DAPI 1 μg/mL if doing immunofluorescence). For immunohistochemistry, Premade Vector ABC mix (Vectastain Elite ABC Reagents, Vector laboratories) was added for 40 minutes. Slides were developed using DAB and DAB-substrate chromogen system (Dako).

### Organoid Wholemount staining

Organoids were single cell dissociated using TrypleE and 15,000 cells were seeded per well in 8-well ibidi chambers (Ibidi, 80826) suitable for confocal imaging. Wholemount staining was performed 7 days post single cell dissociation using standard immunofluorescence wholemount protocol.

### Image analysis

For the whole mount images, ImageJ was used to quantify the number of positive DAPI cells per organoid. The area of interest was circled and the threshold was adjusted. The number of particles were then analysed automatically. The organoid area was also quantified. Three z-stacks per image were analysed and the average was calculated.

### Protein analysis using organoid protein lysates

Organoids were spun down and lysed in RIPA buffer (Thermo Scientific, #89900) with the addition of Halt Phosphatase Inhibitor Cocktail (Thermo Scientific #78420). The Wes™ (Biotechne) automated separation module was used according to manufacturer’s instructions. List of antibodies used is shown in Suppl. Table 4.

### Organoid Competition Assay

Organoids were single cell dissociated using TrypLE and a total of 15,000 cells were seeded per well. One third of organoids in a well were harvested to extract DNA 7 days post single cell dissociation and then 14 and 21 days after. ICE Synthego was used to deconvolute the mixed trace and infer the WT and KO percentages. After the day 7 timepoint, organoids were passaged normally without single cell dissociation.

### Conditioned Media Experiment

Organoids were single cell dissociated using TrypE and a total of 10,000 cells were seeded per well. WT cells were cultured either with PTEN KO or WT conditioned media with the addition of growth factors (EGF, Noggin, FGF1, IGF2, Wnt, Rspo). Images were taken every 3h over 7 days in the Incucyte S3.

### RNA extraction

Frozen PTEN WT and KO organoids were thawed at the same time and expanded for 1 week. They were single cell dissociated and 7 days later RNA was extracted using the Arcturus PicoPure RNA extraction kit (Thermofisher, KIT0204) according to manufacturer’s instructions.

### Bulk RNA-seq library preparation

Library prep was performed using the Illumina Stranded mRNA Prep kit (Illumina, 20040532) according to the manufacturer’s instructions. Samples were submitted for sequencing in the Illumina Novaseq platform with 50 bp paired end reads.

### RNA-seq data analysis

Differential expression analysis was performed using DESeq2. An interaction model was used to identify differentially expressed genes.

Downstream analyses were then performed in R using standard RNA-seq packages clusterProfiler, pathview and msigdbr for gene set enrichment (GSEA) (Subramanian *et al*., 2005; Yu *et al*., 2012; Luo and Brouwer, 2013).

KEGG pathway analysis focused on most highly differentially expressed genes (Fold Change >0.7 or < −0.7). Gene Set Enrichment was performed using Hallmark pathways. Enrichment in Gene Ontology Biological Processes analysis using Gene Ontology online resource based on top 500 differentially expressed genes in the dataset (based on descending p-value).

### qPCR validation

cDNA conversion was performed using ProtoScript II RT standard protocol (NEB, #M0368) and Random Primer Mix (NEB, S1330S).

qPCR was performed using TaqMan Fast Universal PCR Master Mix (Thermofisher, 4352042) and Taqman probes and primers for G0S2 (Thermofisher, Hs00274783_s1), ADH1C (Thermofisher, Hs02383872_s1), NUPR1 (Hs01044304_g1) and GCNT2 (Hs00377334_m1) according to manufacturer’s instructions.

## Author contributions

N.S.P. conceptualisation, formal analysis, funding acquisition, methodology, investigation, validation, visualisation, writing-Original Draft, writing-Review & Editing. S.A. investigation, formal analysis, validation: P.DS. methodology, LM. methodology, S.M. investigation, formal analysis, R.K. formal analysis, software analysis, A.S. investigation, formal analysis, F. P. resources, K.N. resources, A.R. methodology, M.Z. resources, conceptualisation, D. J. W. conceptualisation, funding acquisition, supervision, writing-Original Draft Preparation, Writing-Review & Editing.

## Acknowledgements

We would like to thank the CRUK Cambridge Institute Histopathology, Microscopy, Genomics and Bioinformatics Core Facilities for their help with this paper. Figures were created with Biorender.com.

