## Supplemental Tables for "Efficient genetic editing of human intestinal organoids using ribonucleoprotein-based CRISPR"

### Supplementary information

***Table 1: WENRAFI media composition.*** *Modified from Fujii et al., 2018. Normal organoid maintenance medium.*

| Optimised organoid media (replacement of p38i)-WENRAFI | Stock concentration | Volume | Final concentration |
|---|---|---|---|
| **ADF+++** | pure | 13.08 | |
| **Wnt3a conditioned medium** | pure | 25 ml | 50% |
| **R-spo conditioned medium** | pure | 10 ml | 20% |
| **Primocin (Invivogen #ant-pm-1)** | 50mg/ml (500x) | 100µl | 500 µg/mL |
| **B-27® Supplement (Thermofisher #17504-044)** | 50x | 1000 µl | 1x |
| **Nicotinamide (Sigma #N0636, in water)** | 1 M (100x) | 500 µl | 10 mM |
| **N-Acetylcysteine (Sigma # A9165, in water)** | 500 mM (400x) | 125 µl | 1.25 mM |
| **A3801 (Tocris #2939, in DMSO)** | 5 mM (10,000x) | 5 µl | 500 nM |
| **mEGF (Thermofisher Biosource #PMG8043)** | 100 ng/µl (2,000x) | 25 µl | 50 ng/mL |
| **mNoggin (Peprotech #250-38)** | 100 ng/µl (1,000x) | 50 µl | 100 ng/mL |
| **IGF-1 (Biolegend, 590904)** | 100 ng/µl | 50 µl | 100 ng/mL |
| **FGF-2 (Peprotech, #100-18B)** | 100 ng/µl | 25 µl | 50 ng/mL |
| Total | | 50 mL | |

***Table 2: Guide RNA sequence for guides used in organoids and primer sequence used for screening.***

| **Target** | **Guide sequence** | **F primer sequence** | **R primer sequence** |
|---|---|---|---|
| PTEN | AAAGACTTGAAGGCGTATAC | GGCAGGTGTCAATTTTGGGG | CCTTGGTACACCCAGCGAT |
| ARID1A | CGGTACCCGATGACCATGCA | GCCTTTGTTTATACCCGGCC | CCACTGCCTTTCATCCCATC |

***Table 3: ENAFI media composition.*** *Used for organoid electroporation.*

| **ENAFI (EGF, Noggin, ADF, FGF2, IGF1)** | **Stock concentration** | **Final concentration** | **ENAFI+ Y+Chir (48h before)** | **ENAFI+ Y+Chir+DMSO (24h before and elec day)** |
|---|---|---|---|---|
| **ADF+++** | Pure | | 24010 | 23697.5 |
| **Primocin (Invivogen, #ant-pm-1)** | 50mg/ml (500x) | 500 µg/mL | 50 | 50 |
| **B-27® Supplement (Thermofisher, #17504-044)** | 50x | 1x | 500 | 500 |

| | | | | |
|---|---|---|---|---|
| **Nicotinamide (Sigma #N0636, in water)** | 1 M (100x) | 10 mM | 250 | 250 |
| **N-Acetylcysteine (Sigma # A9165, in water)** | 500 mM (400x) | 1.25 mM | 62.5 | 62.5 |
| **A3801 (Tocris #2939, in DMSO)** | 5 mM (10,000x) | 500 nM | 2.5 | 2.5 |
| **mEGF (Thermofisher Biosource, #PMG8043)** | 100 ng/µl (2,000x) | 50 ng/mL | 12.5 | 12.5 |
| **mNoggin (Peprotech, #250-38)** | 100 ng/µl (1,000x) | 100 ng/mL | 25 | 25 |
| **IGF-1 (Biolegend, 590904)** | 100 ng/µl | 100 ng/mL | 25 | 25 |
| **FGF-2 (Peprotech, #100-18B)** | 100 ng/µl | 50 ng/mL | 12.5 | 12.5 |
| **Y-27632 (STEM Cell Technologies, 73302)** | 10 mM | 10 uM | 25 | 25 |
| **CHIR99021 (Ref)** | 10 mM | 5 uM | 25 | 25 |
| **DMSO** | | 1.25% | | 312.5 |
| Total | | | 25mL | 25mL |

***Table 4: Primary antibodies used for immunohistochemistry (IHC), immunofluorescence (IF) or Wes™.***

| **Protein** | **Supplier** | **Reference** | **Titre** | **Application** |
|---|---|---|---|---|
| **PTEN** | Cell Signaling | #9552 | 1:300 | IHC |
| **MCM2** | BioRad | MCA1859 | 1:300 | IF |
| **Lamin B1** | Cell Signaling | #12586S | 1:2000 | Wes |
| **p-Akt** | Cell Signaling | #4060S | 1:50 | Wes |
| **GAPDH** | Cell Signaling | #5174S | 1:50 | Wes |
| **PTEN** | Cell Signaling | #9552 | 1:100 | Wes |
